# Multiple parameters shape the 3D chromatin structure of single nuclei

**DOI:** 10.1101/2022.01.16.476319

**Authors:** Markus Götz, Olivier Messina, Sergio Espinola, Jean-Bernard Fiche, Marcelo Nollmann

**Author notes:** current address: PicoQuant GmbH, Rudower Chaussee 29, 12489 Berlin, Germany (Markus Götz).

## Abstract

The spatial organization of chromatin at the scale of topologically associating domains (TADs) and below displays large cell-to-cell variations. Up until now, how this heterogeneity in chromatin conformation is shaped by chromatin condensation, TAD insulation, and transcription has remained mostly elusive. Here, we used Hi-M, a multiplexed DNA-FISH imaging technique providing developmental timing and transcriptional status, to show that the emergence of TADs at the ensemble level partially segregates the conformational space explored by single nuclei during the early development of *Drosophila* embryos. Surprisingly, a substantial fraction of nuclei displayed strong insulation even before TADs emerged. Moreover, active transcription within a TAD led to minor changes to the local inter- and intra-TAD chromatin conformation in single nuclei and only weakly affected insulation to the neighboring TAD. Overall, our results indicate that multiple parameters contribute to shaping the chromatin architecture of single nuclei at the TAD scale.

## Introduction

Chromatin in interphase nuclei is organized at multiple levels, including chromosome territories, A/B compartments, and topologically associating domains (TADs) ^1^. TADs, first observed in ensemble-averaged Hi-C contact maps ^2–4^, are sub-megabase genomic regions of preferred contacts and three-dimensional (3D) proximity. In mammalian cells, loop extrusion by the cohesin/CTCF system contributes to TAD formation ^5^. TADs often encapsulate *cis*-regulatory elements (CREs), thereby facilitating interactions between enhancers and promoters that are critical for transcriptional regulation ^6–11^. At the same time, TAD borders may also restrict interactions between CREs located in neighboring TADs ^12–14^. The precise interplay between TADs, enhancer-promoter (EP) contacts, and transcriptional activation is currently under intense study ^15^. Cell type-specific EP contacts have been observed ^16–18^ and direct visualization of EP interactions suggests that sustained physical proximity is necessary for transcription ^19^. On the other hand, other studies suggested that enhancer action may not require loop formation between enhancers and promoters ^20–22^. TADs arise during early stages of development, concomitantly with the activation of zygotic gene expression ^23,24^; however, emergence of TADs seems to be independent of transcription itself in most organisms ^23,25–27^. Remarkably, chromatin structure at the TAD scale is cell-type independent ^28,29^ and does not change upon transcriptional activation ^29^ during early *Drosophila* development. Thus, it is still unclear whether formation of TADs contribute to transcriptional regulation and what is the role played by single-cell heterogeneity.

Several lines of evidence clearly established that chromosome organization is highly heterogeneous between cells. On the one hand, single-cell Hi-C ^30^ revealed that the conformation of individual TADs and loops varies substantially during interphase ^31,32^, along the cell cycle ^33^, and during early development ^34^. On the other hand, imaging-based technologies showed that physical chromatin contacts within and between TADs are rare ^35,36^, and display large cell-to-cell variability ^35–38^. Finally, a high degree of heterogeneity in chromatin organization at the TAD scale was also present in polymer model simulations ^39–41^. Overall, these studies suggest that TADs may exist in the ensemble but not in single cells.

This hypothesis was recently challenged using super-resolution microscopy. Several studies observed that chromosomes folded into ‘nano-compartments’ possibly representing ‘TAD-like domains’ ^42,43^. The condensation of chromatin within nano-domains tends to correlate with epigenetic state ^42^. Borders between nano-compartments appear to be permissible, with a substantial overlap between neighboring regions ^42,44,45^. Consistently, borders between nano-compartments detected in single cells do not necessarily align with ensemble TAD boundaries ^38^. Thus, TAD-like domains exist but display different structural properties between different single cells. How these structural properties relate to single-cell chromatin structures and to transcriptional regulation is currently unclear.

Here, we used Hi-M, a microscopy-based chromosome conformation capture method, to study single-nucleus chromatin organization at the sub-megabase scale before and after emergence of TADs during *Drosophila* embryogenesis. We found a large heterogeneity in single-nucleus chromatin conformations, independent of the presence of a TAD border in the population-average. Remarkably, despite this heterogeneity, chromatin structures of nuclei from different developmental stages segregate in a high-dimensional conformation space. This segregation cannot be assigned to any specific structural parameter. Notably, the single-nucleus chromatin organization of transcriptionally active and inactive nuclei were indistinguishable, both between TADs and inside a TAD. Therefore, chromatin organization is not predictive for the transcriptional state at the single-nucleus level. Finally, the spatial separation of genomic regions from two neighboring TADs was similar for active and inactive nuclei, indicating that physical encapsulation of gene regulatory elements may not be a strict necessity for the spatio-temporal control of transcription during early *Drosophila* embryogenesis.

## Results

### Quantification of single-nucleus chromosome organization heterogeneity during early Drosophila embryogenesis

After fertilization, the fruit fly embryo undergoes thirteen rapid and synchronous nuclear division cycles (nc1 to nc14). During these cycles, nuclei continuously decrease their volumes due to the increase in the number of nuclei and to their migration to a reduced region close to the periphery of the embryo ^46^. Transcription by the zygote is initiated in two waves: the minor wave between nc9-nc13, and the major wave at nc14 ^47–49^. This last stage coincides with the emergence of ensemble TADs ^23^. Thus, early *Drosophila* embryonic development represents an ideal model system to investigate whether heterogeneity in chromosome organization changes with nuclear condensation, with the onset of transcription, and/or with the emergence of TADs.

To address this question, we monitored changes in 3D chromatin organization at the TAD scale between nc11 and nc14 using HiM, a single nucleus multiplexed imaging method that allows for the single nucleus reconstruction of chromatin architecture in whole-mount embryos (Fig. 1a) ^37,50^. We applied Hi-M to a locus displaying two TADs (TAD1 and doc-TAD) (Fig. 1b) annotated using Hi-C data ^23^. The doc-TAD contains three developmental genes (*doc1, doc2*, and *doc3*) ^51^ that are specifically activated at nc14 in the dorsal ectoderm ^29^ (Fig. 1b). Nuclei and barcodes were segmented, localized, and drift-corrected as indicated previously (Figs. S1a-b, ^29,50^), and ensemble Hi-M pairwise distance (ePWD) maps for embryos at nc11/12 and nc14 were constructed by kernel density estimation of the full pairwise distance distributions (Figs. 1b, and S1c-d, Methods). The most notable difference between nc11/12 and nc14 resided in an overall increase in distances between TAD1 and doc-TAD barcodes (Fig. 1c, pink box), and an overall TAD condensation (Fig. S1e, and Fig. 1c, yellow box). These results are consistent with the emergence of ensemble TAD at nc14 ^23^, but do not shed light into the possible changes in structural heterogeneity between nuclear cycles.

**Figure 1.**
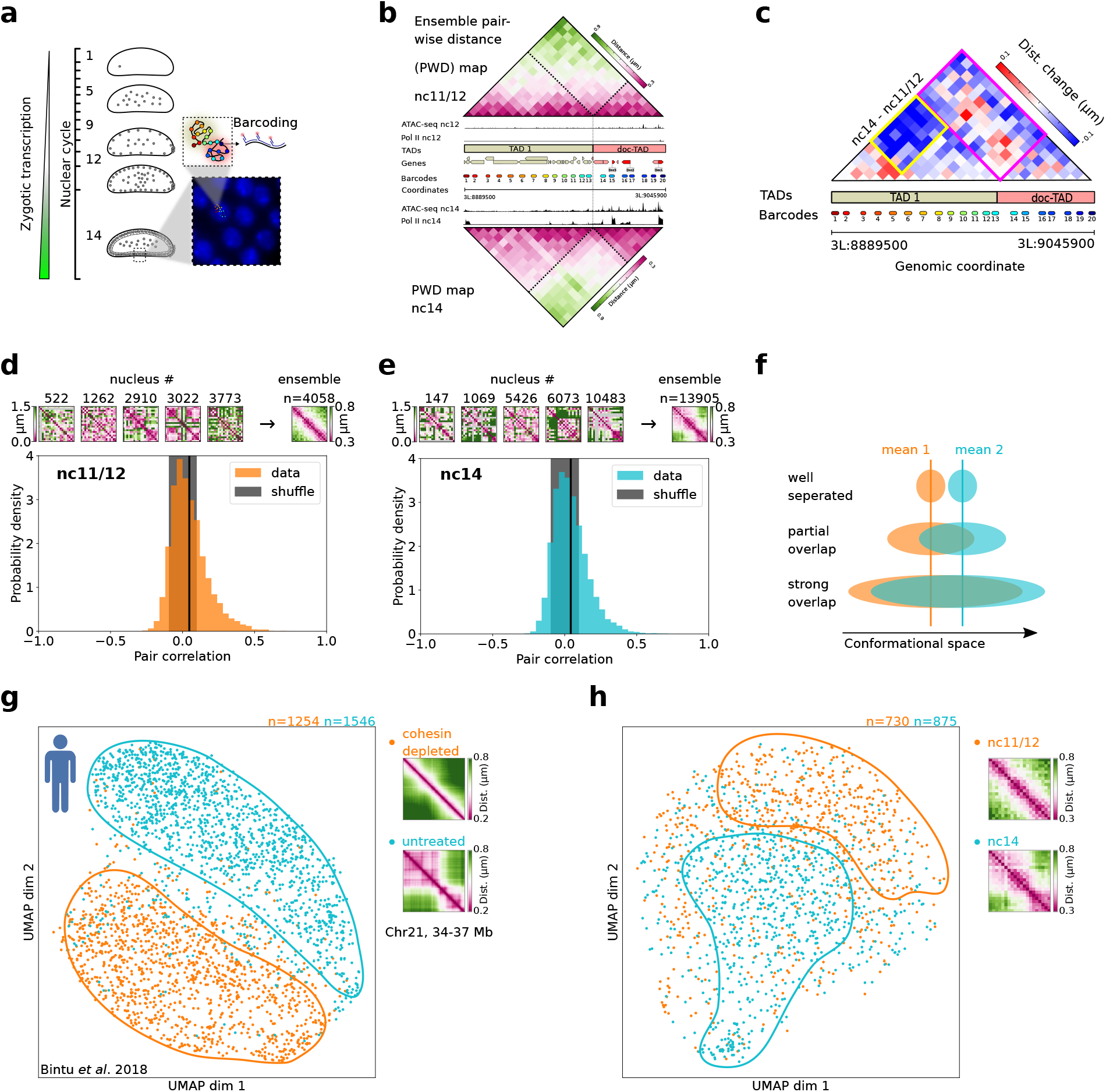
The conformational space explored by chromatin conformations of single nuclei evolves during development. **a**. Scheme of the nuclear positions during the early *Drosophila melanogaster* development as well as the oligopaint-FISH labeling and barcoding strategy. The inset on the bottom right shows DAPI-stained nuclei of a nc14 embryo. The inset on the middle right represents 20 barcodes, each of which comprises 45 oligos with genomic homology. **b**. Extended genomic region (chr3L:8.8895-9.0459 Mb, dm6) around the *doc* locus. Hi-M ensemble pairwise distance maps of nc11/12 and nc14 are shown on the top and bottom, respectively. Tracks of regions with accessible chromatin and with RNA polymerase (ATAC-seq and Pol II ChIP-seq) are shown for both developmental stages. TAD calls from ^23^, and position of genes and barcodes are indicated. Dashed lines in Hi-M maps represent the positions of ensemble TADs from ^23^. **c**. Change in the ensemble pairwise distance maps between nc11/12 and nc14. Pairwise distances that are larger in nc14 than in nc11/12 are shown in red. The pink box highlights the region between TAD1 and doc-TAD, and the yellow box highlights a subTAD within TAD1. **d**. Single-nucleus similarity of the chromatin organization during nc11/12. Top: Five sn pairwise distance maps and the ensemble pairwise distance map. Distances (in µm) are color coded according to the colorbars. Bottom: Histogram of the pair correlation for all pairs of nuclei. A pair correlation of 1 corresponds to identical chromatin organization (apart from a possible constant scaling factor) while a pair correlation of 0 means no correlation between the pairwise distances between two nuclei. The black vertical line indicates the mean pair correlation. The gray area shows the mean +/- standard deviation of the pair correlation for randomly shuffled pairwise distance matrices. **e**. Similar to d, but for nc14. **f**. Scheme of the chromatin conformational space in three hypothetical scenarios: 1. chromatin conformations between two conditions are distinct, leading to their segregation in conformational space. 2. Conformations are partially shared between conditions, leading to an overlap of the occupied conformation space. 3. Conformations are largely the same between two conditions. Note that in all three cases, the ensemble-average of the conformation is the same. **g**. Left: UMAP of the sn chromatin organization in human HCT116 cells in untreated (cyan) and cohesin depleted (orange) conditions. Contours with solid lines highlight regions with a high density of cells from one condition. Data taken from ^38^, locus chr21:34.6Mb-37.1 Mb. Right: Ensemble-average pairwise distance maps for both conditions. **h**. Similar to g, but for data from this study (nuclei of intact *Drosophila* embryos), nc11/12 (orange) and nc14 (cyan).

To characterize variability in chromatin organization during early *Drosophila* development, we estimated the similarity in 3D conformation between different single nuclei. For this, we calculated the correlation between PWD maps of single-nuclei (snPWD) following the method of Conte *et al*. ^41^. We performed this analysis for embryos at nc11/12 and at nc14, before and after the emergence of ensemble TADs. Notably, the PWD maps of individual nuclei displayed a large degree of heterogeneity, with few nuclei exhibiting a PWD map similar to that of the ensemble (Figs. 1d-e, top panels). For both developmental stages, the distribution of pair correlations were broad and centered around zero, indicating that the chromatin conformations of most nuclei were different from each other. Nevertheless, the distribution was skewed towards positive values, indicating that a small proportion of nuclei displayed similar chromatin conformations (Figs. 1d-e, black line indicates median of the distribution). The overall lack of similarity between individual nuclei was observed before and after ensemble TADs emerged (nc11/12 versus nc14, Figs. 1d-e). We note that the lack of similarity between single-nuclei and ePWD maps is perhaps not surprising, as the average of a multi-parametric, widespread distribution tends not to represent any single individual ^52^. Overall, these analyses indicate that chromosome structure heterogeneity is not only present in cultured cells ^35,36,39^ but is also common during early *Drosophila* development, despite the presence of robust mechanisms to control and coordinate gene expression and to synchronize the cell cycle of different nuclei.

### Presence of an ensemble TAD border partially segregates the conformational space explored by single nuclei

Our previous analyses indicate that nc14 and nc11/12 nuclei display distinct ensemble average conformations despite their highly heterogeneous chromatin organizations (Figs. 1c-e). However, these analyses do not reveal the extent to which the conformational spaces explored by single nc14 and nc11/12 nuclei overlap (Fig. 1f): this quantification is needed to estimate whether and how single nuclei structures evolve during development. Thus, we turned to analysis techniques that do not rely on averages but rather on exploring chromatin conformations of single nuclei. The total number of independent dimensions needed to describe the accessible chromatin conformational space can be estimated by the number of degrees of freedom. Even for a small number of barcodes (20), this results in a 54-dimensional space which is inaccessible using conventional plotting methods. Thus, we turned to Uniform Manifold Approximation and Projection (UMAP), an unsupervised, nonlinear dimension-reduction approach previously used to represent single-cell chromatin conformations ^53^. In our implementation, we used UMAP to embed snPWD maps in a 2D space (Methods).

To validate our approach, we applied it to published single-cell data from cultured human cancer cells (HCT116) ^38^. Untreated cells displayed two clearly visible TADs that faded considerably in cohesin-depleted cells (Fig. 1g, right panels). The UMAP embedding of single-cell PWD maps showed two clearly-separated populations highlighted by orange and cyan contours (Fig. 1g, left panel), delimiting a smaller region where single cells were mixed. We note that a small number of untreated single cells (∼3%) localize to the region occupied by cohesin-depleted cells, and vice versa. We observed similar segregation patterns for subsets containing 20 barcodes spanning either a strong or a weak TAD border (Fig. S1f). Thus, UMAP embedding of snPWD maps shows that removal of TAD borders by cohesin depletion dramatically changes the chromatin structure of most single cells at this locus.

To study whether this change also occurred during the natural emergence of TADs during embryonic development, we applied UMAP embedding to snPWD maps at nc11/12 and nc14. Remarkably, we observed that most single nuclei segregated into two distinct populations corresponding to the different nuclear cycles (Fig. 1h). As for human cultured cells, single nuclei occupied extended regions of the UMAP (Figs. 1g-h), consistent with a large degree of heterogeneity in chromosome structure (Figs. 1d-e). Segregation of single-nuclei in two populations was not sensitive to variations of the UMAP hyperparameters (Fig. S1g). Therefore, we conclude that despite a large degree of heterogeneity, the chromatin structures of single nuclei change considerably during the early cycles of *Drosophila* embryogenesis and occupy distinct conformational spaces.

### TAD condensation is not critical to distinguish between single nuclei conformations

Next, we implemented several analyses to investigate the factors influencing the segregation of single nuclei within the UMAP conformational space. First, we constructed average PWD matrices for a selection of single nuclei located at the four cardinal directions of the UMAP (west, east, south, north) (Fig. 2a). Matrices from west and east nuclei were similar to the ePWD map (Fig. 1b). Instead, south and north maps displayed more marked, complementary differences that can be mapped to the relative positions of nc11/12 and nc14 nuclei within the UMAP space (Fig. 1h), and that mirrors the difference between nc11/12 and nc14 ensemble PWD maps (Fig. 1c). We conclude that in this representation the second UMAP dimension encodes the most important differences between the structures of nc11/12 and nc14 nuclei.

**Figure 2.**
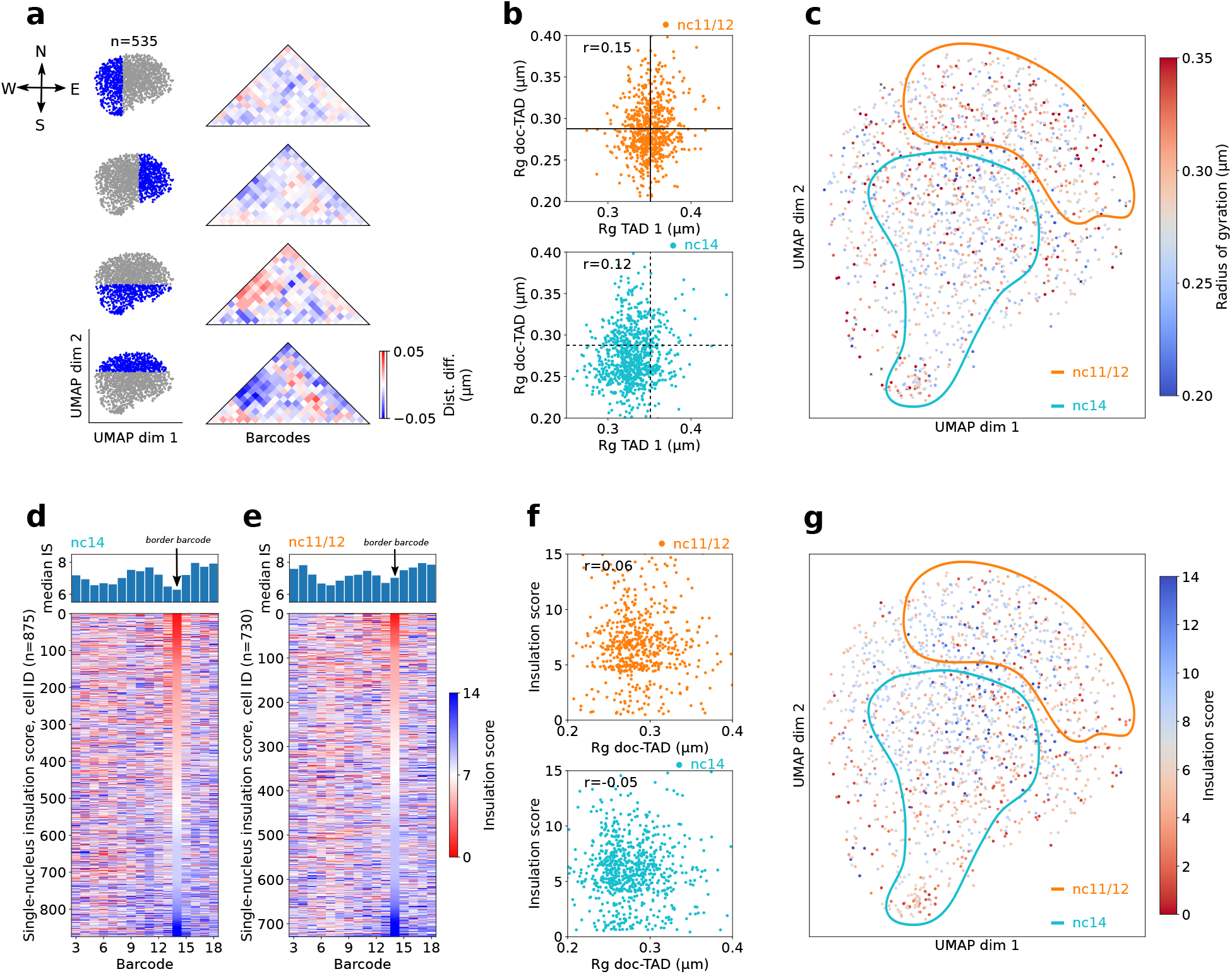
Segregation of nuclei populations in the UMAP space is due to a combination of multiple architectural features. **a**. Interpretation of the first two UMAP dimensions. Left: Selection of one-third of the nuclei in the UMAP (indicated by blue color). Right: The difference of the ensemble pairwise distance map of the selection and all cells in the UMAP. **b**. Top: scatter plot of the correlation between TAD volumes, as measured by the radius of gyration, for doc-TAD and TAD1 in nc11/12 nuclei. Each point represents a single nucleus. The horizontal and vertical lines indicate the mean of the distribution. *r* is the Pearson correlation coefficient. Bottom: similar to the top, but for nc14. The dashed lines indicate the mean of nc11/12. **c**. Same UMAP as in Fig. 1h, color-coded by the radius of gyration of the doc-TAD. Blue and red correspond to small and large radii, respectively. **d**. Top: Ensemble-average (median) profile of the insulation score for nc14. The black arrow indicates the border barcode. Bottom: Single-nucleus insulation score. Each line corresponds to the insulation score profile of an individual nucleus, with values color-coded from red (small insulation score, i.e. strong insulation) to blue (large insulation score, weak insulation). The color scale is the same as in e. Nuclei are sorted according to their insulation score at the border barcode. **e**. Similar to d, but for nc11/12 nuclei. **f**. Top: scatter plot of insulation score at the border barcode versus radius of gyration of the doc-TAD in nc11/12. Each point corresponds to a single nucleus. *r* is the Pearson correlation coefficient. Bottom: similar to the top, but for nc14. **g**. Same UMAP as in Fig. 1h, color-coded by the insulation score at the border barcode. Blue and red colors correspond to small and large insulation scores (i.e. strong and weak insulation), respectively.

Second, we explored the role of TAD condensation by measuring the radius of gyration (R_g_) of the two TADs in single nuclei as a proxy for TAD volume. Adjacent TAD volumes were highly heterogeneous and uncorrelated (Pearson coefficient ∼0.1), but decreased from nc11/12 to nc14 (Fig. 2b). This latter observation raises the possibility that changes in TAD volumes may contribute to the segregation of the two distinct UMAP populations. To explore this hypothesis, we color-coded each nucleus in the UMAP by the radius of gyration of the doc-TAD (Fig. 2c). Volumes in nuclei within the nc14 population tended to exhibit smaller volumes (R_g_<0.27μm), while the opposite was observed in the nc11/12 population (R_g_>0.27μm). However, nuclei in both regions displayed extended and compact TAD volumes and no clear trend of the radius of gyration along either UMAP dimension was discernible. Thus, TAD volume contributes but is not the only factor separating single nuclei within the UMAP space.

### Single nuclei displaying insulated TADs are common before the emergence of ensemble TADs

Next, we investigated the role of TAD insulation in the segregation of single nuclei conformations. For this, we first calculated the ensemble and single nuclei insulation scores (IS) of nc14 nuclei (Fig. 2d, top panel) following the method developed by Crane *et al*. ^54^. Low insulation scores represent regions with strong borders (see Methods). As expected, the median IS profile for nc14 nuclei showed a dip at the barcode located at the border between TAD1 and doc-TAD (hereafter *border barcode*). The corresponding dip in the median IS profile for nc11/12 was instead less pronounced (Fig. 2e, top panel), consistent with the emergence of the ensemble TAD border at nc14 in this locus (Figs. 1b-c). We explored the variability of this border by stacking the single-nuclei IS profiles, sorted by their IS score at the border barcode (Fig. 2d,e, bottom panels). We quantified the proportion of nuclei displaying a strong insulation between TADs by using a single-nucleus IS cutoff of 3.5. Notably, at nc14 only a small proportion of single nuclei (∼14% with IS < 3.5) displayed insulation at the border barcode (i.e., the ensemble TAD border), while many nuclei exhibited low insulation scores at other genomic locations. Similar results were obtained for other IS cutoffs (Fig. S2a). Thus, we conclude that for nc14 embryos the border barcode is not always insulated but rather represents the region displaying the most preferred insulation, consistent with results in human cultured cells ^38^.

As ensemble TADs emerge at nc14, we wondered whether and to what extent single nuclei in previous nuclear cycles were already insulated at the border barcode. As expected, the ensemble IS at the border barcode was higher in nc11/12 than in nc14 nuclei (Fig. 2e, top panel). However, we were surprised by the frequency of single nc11/12 nuclei displaying insulation at the border barcode. This frequency was smaller (10% with IS < 3.5), yet comparable to that observed in nc14 embryos, indicating that single nuclei displaying insulated TADs also exist in early developmental cycles before the emergence of ensemble TADs. Notably, TAD condensation does not seem to define the level of TAD insulation, as insulation scores and TAD volumes were uncorrelated both in nc11/12 and nc14 embryos (Fig. 2f, S2b).

Finally, we explored whether the presence of a TAD border was sufficient to split cells between UMAP populations by color-coding each single nuclei in the UMAP by the insulation score at the border barcode (Figs. 2g and S2c). We observed a wide distribution of insulation scores for both UMAP populations and no strong correlation with any of the UMAP dimensions. We observed a similar broad distribution of insulation scores for human HCT116 cells (Fig. S2c). Remarkably, nuclei with high insulation scores (i.e. low insulation) were common within the nc14 UMAP population (Fig. 1g). Similarly, single cells with high IS were common in untreated human HCT116 cells (Fig. S2c). Importantly, single nuclei in nc11/12 commonly displayed low IS values (i.e. high insulation), an observation that is mirrored by cohesin depleted HCT116 single cells (Fig. S2c). In summary, only a minority of nuclei exhibit a strong border between TAD1 and doc-TAD in nc14 embryos, and this population of insulated TAD is already present at earlier stages of development that do not display discernable ensemble TADs.

### Transcriptionally active and inactive nuclei explore similar regions of the UMAP space

Naturally, we wondered whether transcription contributed to the single-nucleus organization of chromatin at the *doc* locus. To address this question, we imaged DNA organization by Hi-M together with the detection of *doc1* expressing nuclei by RNA-FISH in the same embryos (Fig. 3a). The presence of *doc1* nuclear transcription hotspots was used to label each single nuclei in the embryo as *doc1* active or inactive (Fig. 3a). In a first attempt to correlate TAD organization and transcription, we calculated the median intra- and inter-TAD distances for individual active and inactive nuclei. As expected, intra-TAD distances were smaller than inter-TAD distances (Fig. 3b, Methods). Interestingly, both intra- and inter-TAD distances were higher for transcriptionally active nuclei. Similar trends were observed for the difference PWD map and for the distribution of doc-TAD volumes in active and inactive nuclei (Figs. S3a-b). Overall, these results indicate that the doc-TAD was −on average− slightly more decondensed and more segregated from TAD1 in actively transcribing nuclei than in inactive nuclei.

**Figure 3.**
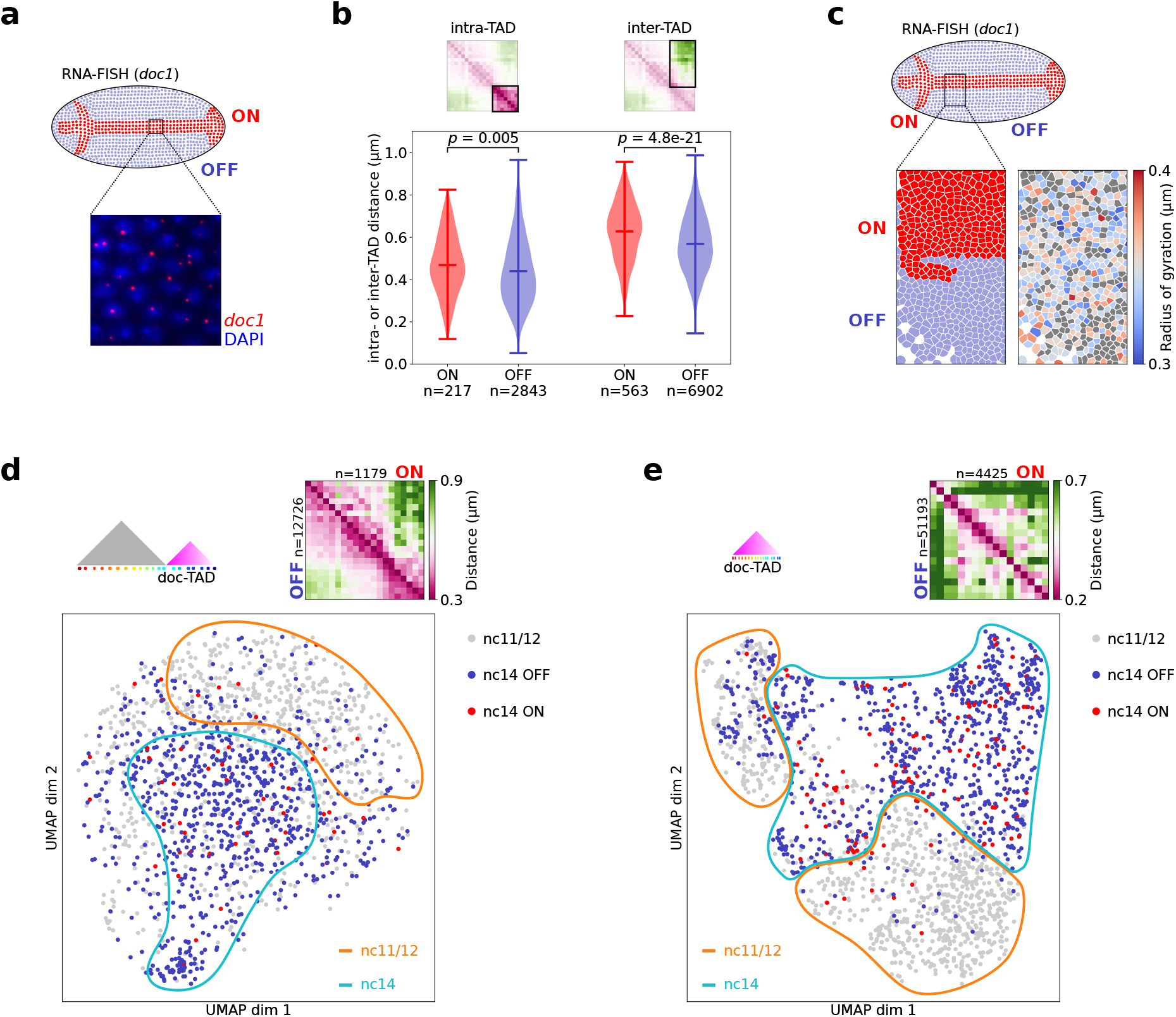
Transcriptionally active and inactive nuclei explore a similar conformational space. **a**. Top: scheme of a *Drosophila* embryo in dorsal orientation. Nuclei transcribing *doc1* are indicated in red (“ON”), nuclei not transcribing *doc1* in blue (“OFF”). Bottom: DAPI-stained nuclei (blue) with *doc1* RNA-FISH spots indicating nascent transcripts (red). **b**. Intra- and inter-TAD distance distributions shown as violin plots. For each nucleus, the median of the pairwise distances highlighted in the distance map on the top is calculated. Distances for ON nuclei are shown in red, distances for OFF nuclei in blue. Markers indicate the mean and extreme values of the distribution. **c**. Bottom left: Cell masks after DAPI image segmentation from a Hi-M experiment. The color indicates the transcriptional state. Bottom right: The same masks as on the left are color-coded by the radius of gyration of the doc-TAD. Nuclei for which no radius of gyration could be calculated are in gray. **d**. Top: Ensemble-average pairwise distance map for ON (top right half of the matrix) and OFF nuclei (bottom left half) for the low-resolution Hi-M library. Bottom: Same UMAP as in Fig. 1h, color-coded by nuclear cycle and transcriptional state. nc11/12 nuclei are in gray, nc14 OFF in blue and nc14 ON in red. **e**. Similar to b, but for the high-resolution Hi-M library, that spans the doc-TAD with higher resolution.

To determine whether this trend was also visible in single nuclei, we calculated the radius of gyration of the *doc* TAD in each single nucleus and displayed it along the *doc1* activation pattern (Fig. 3c). We observed that the volume of the *doc* TAD did not strongly correlate with the transcriptional status of single nuclei. Notably, nuclei with large and small TAD volumes were observed both in the active and inactive patterns (Fig. 3c, bottom panels). Thus, TAD volume seems to only poorly distinguish between single nuclei with different transcriptional states.

Different chromatin structures within TADs may give rise to similar TAD volumes, thus we turned to UMAP embedding to further explore the possible role of transcription in the 3D chromatin structure of single nuclei. For this, we embedded single nc11/12 and nc14 nuclei together, and color coded them according to their transcriptional status (Fig. 3d). *Doc1* is expressed at nc14, thus most active cells appeared within the region of the UMAP enriched in nc14 nuclei (Fig. 3d, cyan shape). Remarkably, active and inactive cells were homogeneously distributed across this region, consistent with transcription not playing a key role in determining the overall 3D conformation of single nuclei.

The genomic coverage of the oligopaint library used in these experiments was sufficient to visualize TADs in this locus, but not to detect specific regulatory interactions within the doc-TAD (Fig. 3d). Thus, we performed a similar analysis using a published dataset with a genomic coverage that enabled the detection of cis-regulatory interactions (Fig. S3c) ^29^. Nc11/12 and nc14 cells occupied different regions of the UMAP embedding (Fig. 3e), consistent with the results obtained with the lower coverage oligopaint library (Fig. 1h). In addition, *doc1* active and inactive nuclei also intermingled within the nc14 pattern and did not segregate from each other. Consistent with these findings, TAD volumes were similar for active and inactive nuclei for the higher resolution library (Fig. S3d). All in all, these results suggest that the chromatin organization of active and inactive nuclei are indistinguishable at the ensemble and single nucleus levels.

### Single nucleus TAD insulation and intermingling are independent of transcriptional activity

Several lines of evidence suggest that TAD borders play a role in insulating enhancer-promoter interactions between neighboring TADs, however this role is currently under intense debate ^15^. Thus, we wondered whether insulation at the single cell level affected the expression of *doc1*. For this, we plotted the single nucleus IS profiles ranked by IS at the border barcode together with transcriptional status (Fig. 4a). Notably, we did not observe a correlation between the transcriptional state of single nuclei and their insulation score (Fig. 4a). In fact, both active and inactive nuclei displayed low insulation scores (i.e. strong TAD border), and conversely in many active nuclei we failed to observe a clear single-nucleus boundary at the border barcode. These conclusions are supported by the ensemble IS profiles for active and inactive cells (Fig. 4b) which show that active cells are similarly insulated than inactive cells.

**Figure 4.**
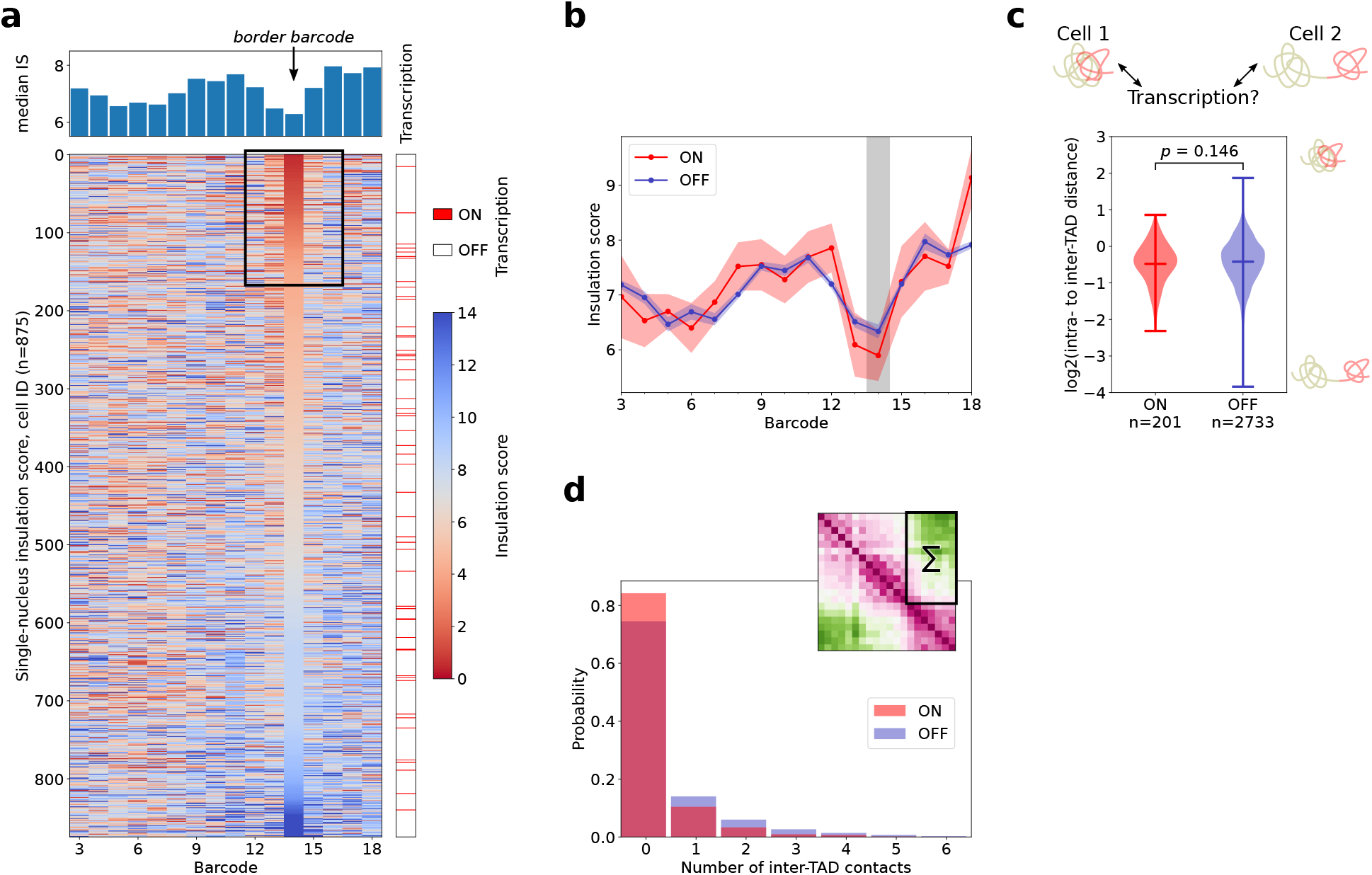
Single-nucleus TAD insulation and intermingling with a neighboring TAD are independent of transcriptional activity. **a**. Top: Ensemble-average (median) profile of the insulation score for nc14 embryos. Bottom: Single-nucleus insulation score. Each line corresponds to the insulation score profile of an individual nucleus, with values color-coded from red (small insulation score, i.e. strong insulation) to blue (large insulation score, i.e. weak insulation). Nuclei are sorted according to their insulation score at the TAD border. The right-most lane indicates the transcriptional state of each nucleus, with active nuclei marked red. The black frame highlights the ∼20% of nuclei with a high insulation (i.e. a low insulation score) at the ensemble TAD border. **b**. Ensemble-average profile of the insulation score for active (red) and inactive (blue) nc14 nuclei. Circles indicate the median IS for each barcode. The red and blue shaded bands represent the uncertainty as estimated by bootstrapping. The vertical grey bar indicates the border barcode. **c**. Demixing score of TAD 1 and doc-TAD for active (red) and inactive (blue) nuclei. Large values indicate a stronger intermingling of the two TADs as shown by the scheme to the right. Markers indicate the mean and extreme values of the distribution. **d**. Histogram of the number of inter-TAD contacts between TAD 1 and doc-TAD. For each nucleus, the sum of contacts in the region highlighted in the distance map on the top is calculated. Active nuclei in red, inactive nuclei in blue.

To test whether proximity of TAD1 enhancers to the *doc1* promoter influenced its transcription, we calculated the intermingling between TADs for active and inactive cells. TAD interminging was estimated by the demixing score, calculated by measuring the ratio between intra- to inter-TAD distances (see Methods). Notably, the distributions of demixing scores were very similar for both states of transcription (Fig. 4c). This result is inconsistent with physical proximity between TAD1 enhancers and the *doc1* promoter playing an important role in its transcriptional activation. As PWD distributions are broad, changes in short-range distances, i.e., a contact between loci, might be overlooked when only considering distances ^55^. Therefore, we binarized the single nucleus PWD maps using a contact threshold of 0.25 µm and tested whether the transcriptional state of single nuclei was linked to a change in the number of contacts between the two TADs (Fig. 4d). In fact, the distributions in the number of inter-TAD contacts were very similar for active and inactive nuclei, supporting the conclusions from our previous analysis using demixing scores.

All in all, these analyses show that at this locus TADs are insulated in a small proportion of nuclei, that insulation is not correlated with transcriptional activation at the single nucleus level, and that TAD intermingling is as common in active as in inactive nuclei. Thus, we conclude that transcription does not seem to play a key role in the structure of single nuclei as measured by TAD insulation and intermingling.

## Discussion

Population-average TADs require the presence of CTCF and cohesin in mammals ^38,56–58^, and can first be detected at the zygotic genome activation step in multiple species ^23,24,59–61^. In this study, we used an imaging-based method that simultaneously provides developmental timing, transcriptional status, and snapshots of chromatin conformations in single nuclei during this developmental transition. This imaging method, combined with state-of-the-art single-cell analysis techniques, provides new insights into the roles that TAD insulation, TAD condensation and transcription play in shaping chromatin structure.

Chromatin organization at the TAD scale was previously shown to be highly variable both by FISH ^35,36,38,39^, and by polymer modeling simulations ^5,40,41,62^. These models showed that chromatin forms globular TAD-like structures in the presence of attractive interactions between monomers in a TAD ^40^, or in the presence of loop extrusion by cohesin ^63,64^. In contrast, chromatin behaves as a random-coil polymer in the absence of cohesin or in absence of monomer-monomer interactions ^40,63,64^. Notably, our results show that chromatin is similarly heterogeneous in presence or absence of ensemble TADs, suggesting that intra-TAD interactions do not play a major role in constraining the degree of variability of chromatin conformations. A reasonable explanation for this behavior, compatible with simulation and imaging data ^35,36,40^, is that intra-TAD interactions are transient and highly variable between cells, likely reflecting the binding dynamics of transcription factors and architectural proteins ^65–68^ and their cell-to-cell heterogeneity, as well as chromatin conformational dynamics.

Notably, despite this heterogeneity, the chromatin conformations of single nuclei before and after the emergence of ensemble TADs were distinct enough to occupy different regions in the 2D UMAP embedding of the conformational hyperspace. We observed similar results in datasets from untreated and cohesin-depleted human cells. Thus, tracing the path of chromatin in a single nucleus at the TAD-scale and with relatively sparse sampling (20-50 barcodes) is sufficient to predict, with reasonable confidence, the presence of mechanisms leading to TAD formation. We insist, however, on the probabilistic nature of this prediction, given that (1) single nuclei conformations may fall, with low probability, into regions of the UMAP space displaying overlapping conformations; and (2) the regions of the UMAP space attributed to each population are not unequivocal.

These results suggested that structural parameters, such as TAD insulation or condensation, or transcriptional activity may be the main factors segregating single nuclei in the UMAP space. Surprisingly, this does not appear to be the case, as none of these factors played predominant roles in the segregation of single conformations in the UMAP space. For instance, while TADs tended to be more condensed in nc14 embryos, an important proportion of nc11/12 embryos also exhibited TADs condensed to similar levels. Surprisingly, only a fraction of nuclei displayed strong insulation between TAD1 and the doc-TAD in nc14 embryos, and a smaller but significant fraction of nuclei exhibited strong insulation in nc11/12 embryos. These results are consistent with Hi-C and imaging studies where TAD borders in *Drosophila* were shown to be variable between single nuclei ^31,35^. In *Drosophila*, TAD boundaries are mainly occupied by architectural factors ^13,14^, and their appearance requires binding of pioneering factors ^23^. In this context, our results suggest that occupation of TAD borders: (1) is stochastic, possibly due to the binding kinetics of architectural proteins and of components of the transcriptional machinery; (2) gradually increases during early development as these factors become more abundant. All in all, our results suggest that neither TAD condensation, nor TAD insulation can be used to predict whether a nucleus belongs to nc14 or nc11/12 embryos. We conclude that, instead of a single structural parameter, multiple combinatorial contributions from several structural properties may be required to assign single chromatin conformations to specific developmental stages with high confidence. This is consistent with a recent study showing that multiple, spatially-distributed structural features are required to predict transcriptional activation in the BX-C domain during late Drosophila embryogenesis ^69^.

Cell-to-cell variations in the gene activation of genes within a TAD, either arising from controlled variations in transcriptional programs between cell-types or from stochasticity in transcription ^70^, could arguably contribute to the spatial distribution of single conformations within the UMAP space. Consistent with this idea and with previous evidence ^35,42^ active nuclei showed on average a slightly more decondensed chromatin architecture than inactive nuclei. However, despite these small differences, transcriptionally active and inactive nuclei explored overlapping conformational spaces and did not segregate from each other in the UMAP space. These results are consistent with recent studies showing that, during *Drosophila* development, the population-averaged chromatin architectures of cells from different presumptive tissues display only overall small differences ^28,29^. Thus, we conclude that transcription does not seem to play a dominant role in determining the chromatin structure of single nuclei at least during these early stages of *Drosophila* development.

The single nucleus snapshots of chromatin architecture occupied preferential regions in the UMAP space that depended on the developmental stage. How much of this conformational space can be explored by an individual nucleus during a biologically relevant time-scale, i.e., one cell cycle? The succession of nuclear division cycles is fast during early development of *Drosophila*, with cell cycles lasting 10-12 minutes for nc11 and nc12 and at least 65 minutes for nc14 ^71^. Existing experimental approaches that dynamically track genomic loci ^19,21,72^ are currently not able to address the question of TAD-scale chromatin mobility directly as technological limitations in live-cell labeling so far have prevented dynamic visualization of more than two loci at the same time. Future studies that simultaneously track more than two genomic regions *in vivo* at high genomic (<10kb) and optical (<150nm) resolutions will be needed to determine whether chromatin within TADs dynamically explore their conformational space during interphase. Nevertheless, polymer physics predicts that a spatial neighborhood of 150-300nm (roughly corresponding to a genomic region of 100kb) can be extensively explored by a locus in 2-5 minutes ^73,74^. This suggests that, despite short cell cycle times during early *Drosophila* development, a genomic region spanning a few TADs (as that investigated here) should be able to cover a substantial part of the conformational space outlined by single nucleus snapshots, leading to ergodicity at these short spatial scales.

Studies in mammalian cells have shown a non-zero probability for TAD-like domain boundaries at any locus ^38^ and a heterogeneity in the intermingling of two regions that are separated by a TAD border in the ensemble-average ^43,44^. Our model system enabled the investigation of single nucleus variation in TAD insulation in intact embryos, where development progresses in a highly synchronous and orchestrated fashion in contrast to cell cultures. Thus, we expected the contribution of external sources of perturbation (e.g. differences in cell cycle stage or transcriptional activation within well-defined presumptive tissues) to be reduced. Nevertheless, we still observed that insulation between adjacent TADs is highly heterogeneous at the single nucleus level. Additionally, we found that intermingling of the doc-TAD with the upstream TAD is independent of *doc1* activation. This lack of a strong insulation effect suggests that tissue- and time-specific enhancers carrying the right composition of transcription-activating factors, together with promoters bound by compatible sets of factors, are the main regulators of gene expression in this system. In this scenario, other nearby enhancers would only play a minor role, due to lack of compatible activating factors, even when they could be in spatial proximity to the promoter. Alternatively or in addition, a carefully-balanced non-linear relationship between enhancer-promoter contact frequency and transcriptional output could also explain the similar single nucleus insulation profile of active and inactive nuclei. Such non-linear models have been proposed by several groups recently ^75–77^. These non-linear models have in common that gene activation is modeled as a multi-step process that makes repeated interactions between enhancer and promoter necessary before gene activation is achieved. Thus, EP contacts spanning a population-average TAD border could occur, but with a frequency that is not sufficient to effectively trigger transcription.

In the case of the *doc* genes, multiple enhancer elements (validated or putative) are distributed throughout the doc-TAD ^29^. As the number of enhancers vastly outnumbers the number of genes ^78^, regulation of a gene by multiple enhancers seems to be the rule, not the exception. The classical gene regulation model assumes that stable, long-lasting EP contacts need to be formed, thus restricting the possibility for alternative chromatin configurations. By contrast, a highly flexible chromatin organization could be an advantage, as different enhancers could physically access the promoter region, thereby offering a way to ensure phenotypic robustness as discussed for mammalian organisms ^79^. This robustness would be especially important during early development when cells need to follow precise spatiotemporally gene-expression programs. It is interesting to note that a recent deep-learning analysis of chromatin tracing data suggest that chromatin structures linked to the transcriptional state of a gene are broadly distributed across the gene’s regulatory domain and that individual enhancer-promoter interactions don’t play a major role in defining the transcriptional activity of a gene ^69^.

Overall, our analysis of single-nucleus microscopy-based chromosome conformation capture data is compatible with a model of flexible chromatin organization that serves as a scaffold with enhancer-bound transcription-activating factors encoding the logic that integrates multiple, potentially short-lived, interactions that can extend beyond domain borders defined from ensemble-averaged experiments.

## Acknowledgements

This project was funded by the European Union’s Horizon 2020 Research and Innovation Program (Grant ID 724429) (M.N.). We acknowledge the Bettencourt-Schueller Foundation for their prize ‘Coup d’élan pour la recherche Française’, the France-BioImaging infrastructure supported by the French National Research Agency (grant ID ANR-10-INBS-04, ‘‘Investments for the Future’’), and the Drosophila facility (BioCampus Montpellier, CNRS, INSERM, Univ Montpellier, Montpellier, France). M.G. was funded by the Deutsche Forschungsgemeinschaft (DFG, German Research Foundation) - project ID 431471305. O.M. is supported by an FRM PhD fellowship.

## Data and code

The data used in this manuscript was uploaded to Zenodo (10.5281/zenodo.5815196). The code used in this manuscript is accessible at: https://github.com/NollmannLab/Goetz_etal

## Methods

### Drosophila embryo collection

Embryos from fly stocks (Oregon-R w^1118^) were collected and fixed as previously described ^29^.

### Hi-M libraries

Oligopaint-FISH libraries used in this study have been previously described ^29^. In brief, the standard-resolution library consisted of 20 barcodes and covered two adjacent TADs (3L:8889500..9045900, Release 6 reference genome assembly for *Drosophila melanogaster*), including the *doc* locus. The high-resolution library comprised 17 barcodes in the doc-TAD (3L:8981462..9045820).

### RNA-FISH and hybridization of the primary Hi-M library

The *in situ* hybridization protocol for RNA detection of *doc1* and the protocol for hybridization of the primary Hi-M library are described elsewhere ^29,50^.

### Acquisition of Hi-M datasets

Experiments were performed on a homemade, wide-field epifluorescence microscope described previously ^29^. Protocols for attachment of embryos to the coverslip and for the sequential labeling and imaging were also previously reported ^29^.

### Image processing

The acquired dataset consisted of image stacks with 2048×2048 pixels and 60 slices (voxel size 0.106×0.106×0.250 µm^3^). Raw images supplied by the camera were in DCIMG format and were converted to TIFF using proprietary software from Hamamatsu. The TIFF images were then deconvolved using Huygens Professional v.20.04 (Scientific Volume Imaging, https://svi.nl). Further analysis was done using a homemade software pipeline written in python ^29^. First, images were z-projected using sum (DAPI channel) or maximum intensity (barcodes, fiducials) projection. Then, for each hybridization round, the image of the fiducial channel was aligned to the reference fiducial image in a two-step process: 1) Global alignment by cross-correlation of the two images split in 8×8 non-overlapping blocks and averaging over the translation offset of all 64 blocks. 2) Local alignment in 3D of volumes each containing a single nucleus by cross-correlation.

Barcodes were segmented using a neural network (*stardist*) ^80^ specifically trained for the detection of 3D diffraction limited spots produced by our microscope. To extract the position of the barcode with sub-pixel accuracy, a subsequent 3D Gaussian fit of the regions segmented by *stardist* was performed with Big-FISH (https://github.com/fish-quant/big-fish, ^81^). Barcode localizations with intensities lower than 1.5 times that of the background were filtered out.

Nuclei were segmented from projected DAPI images using *stardist* ^*80*^ with a neural network trained for detection of nuclei from *Drosophila* embryos under our imaging conditions. Barcodes were then attributed to single nuclei by using the XY coordinates of the barcodes and the DAPI masks of the nuclei. Finally, pairwise distance matrices were calculated for each single nucleus.

The transcriptional state of the nuclei was attributed by manually drawing polygons over the nuclei displaying a pattern of active transcription.

### Ensemble-average pairwise distance map

The ensemble pairwise distance map was calculated from the first maximum of the kernel density estimation for each pairwise distance distribution (Gaussian kernel, bandwidth 0.25 pixel, excluding pairwise distances larger than 4.0 µm), which is a robust approximation for the mode of the pairwise distance distribution.

In case of the human HCT116 cell line data ^38^, the median of the pairwise distance distribution was used to calculate the ensemble pairwise distance map, following the approach of the original paper.

### Radius of gyration

The radius of gyration was calculated from the pairwise distances using 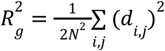, where *d*_*i, j*_ is the pairwise distance between barcode *i* and *j*. The number of pairwise distances (*N*^*2*^) is adjusted in case a pairwise distance was not detected for a given cell. Pairwise distances above a threshold (1.0 µm) were set to NaN to remove outliers that would bias the radius of gyration towards large values. Cells with less than one third of all pairwise distances detected were excluded from analysis.

Calculation of the radius of gyration is based on the standard-resolution Hi-M library except for Fig. 3c, which uses the high-resolution Hi-M data as a slightly higher fraction of nuclei could be used to calculate the radius of gyration for this data.

### Pair correlation

The similarity of the chromatin organization between single nuclei was calculated by the pair correlation for all possible pairs of nuclei, following ^41^.

In more detail, nuclei with a detection of less than 13 out of 20 barcodes (low-resolution library) or 11 out of 17 barcodes (high-resolution library) were excluded from analysis. The upper triangle of the sn pairwise distance maps was flattened to yield a vectorized representation. The Pearson correlation coefficient for all pairs of vectorized distance maps was calculated, using the reciprocal distance to be more sensitive to changes in small distances. Pairwise distances that were not detected in both corresponding nuclei were masked for the calculation of the correlation coefficient. A given pair of nuclei was skipped when the number of the overlapping detected pairwise distances is smaller than half of the full number of pairwise distances.

As a reference, the pair correlation of randomly shuffled pairwise distance maps was calculated. For this, for each pair one of the pairwise distance maps was reordered randomly ten times and the Pearson correlation coefficient was calculated as described above for each of the ten randomizations.

### UMAP embedding

The number of nuclei for different developmental time points (nc11/12 vs nc14) or different treatment (wt vs auxin treated) were roughly matched by adjusting the cutoff for excluding nuclei from analysis based on the number of detected barcodes. For the low-resolution library, nuclei were excluded when more than 7, or 6, out of 20 barcodes were not detected (nc11/12, or nc14, respectively). For the high-resolution library, nuclei were excluded when more than 5, or 2, out of 17 barcodes were not detected (nc11/12, or nc14, respectively). For the human HCT116 cell line data, nuclei were excluded when more than 5, or 1, out of 83 barcodes were not detected (auxin-treated, or wt, respectively).

For the remaining nuclei, single nucleus pairwise distances not detected were imputed with the corresponding value from the ensemble pairwise distance map. Then, the single nucleus pairwise distance maps were vectorized (see “pair correlation” above) and the different categories (developmental time points or treatments) were concatenated. Dimension reduction to two dimensions was achieved by unsupervised embedding, using the python implementation of “Uniform Manifold Approximation and Projection” (UMAP) ^82^. Parameters were: n_neighbors=50, min_dist=0.1, n_epochs=500, metric=“canberra”.

### Insulation score

If indicated, the same selection of nuclei and imputation of missing sn pairwise distances as for the UMAP embedding was performed. Otherwise the raw data was used.

To calculate the sn insulation score, we followed an approach similar to the one used for bulk Hi-C contact maps ^54^. In short, a square of 2-by-2 barcodes is moved parallel to the diagonal of the pairwise distance map and the inverse of the pairwise distances in this square are summed. This yields a profile of the insulation score per nucleus with lower insulation score values corresponding to a higher insulation.

### Intra- and inter-TAD distances and demixing score

To get average intra- and inter-TAD distances for each nucleus, the median of all pairwise distances in the doc-TAD or between TAD 1 and doc-TAD were calculated. Distances above a threshold (1.0 µm) were excluded and medians were calculated only when at least 3 intra- or inter-TAD distances were detected for a nucleus.

The demixing score is calculated per nucleus as the log2 ratio of the median intra- and inter-TAD distances.

### Inter-TAD contacts and 3-way interactions

Two barcodes were considered in contact when their distance was less than 250 nm. The number of inter-TAD contacts per nucleus was obtained by counting the number of contacts in the inter-TAD region in the contact map.

We calculated the 3-way contact probability as the number of barcode pairs within 250 nm to any third barcode, normalized by the number of nuclei containing all three barcodes. This approach thus highlights barcode pairs that are frequently involved in 3-way interactions.

### Statistics

The *p*-values in Fig. 3b, 4c, and S1e were calculated by a two-sided Welch’s *t*-test.

## Figure Legends

**Supplementary Figure 1.**
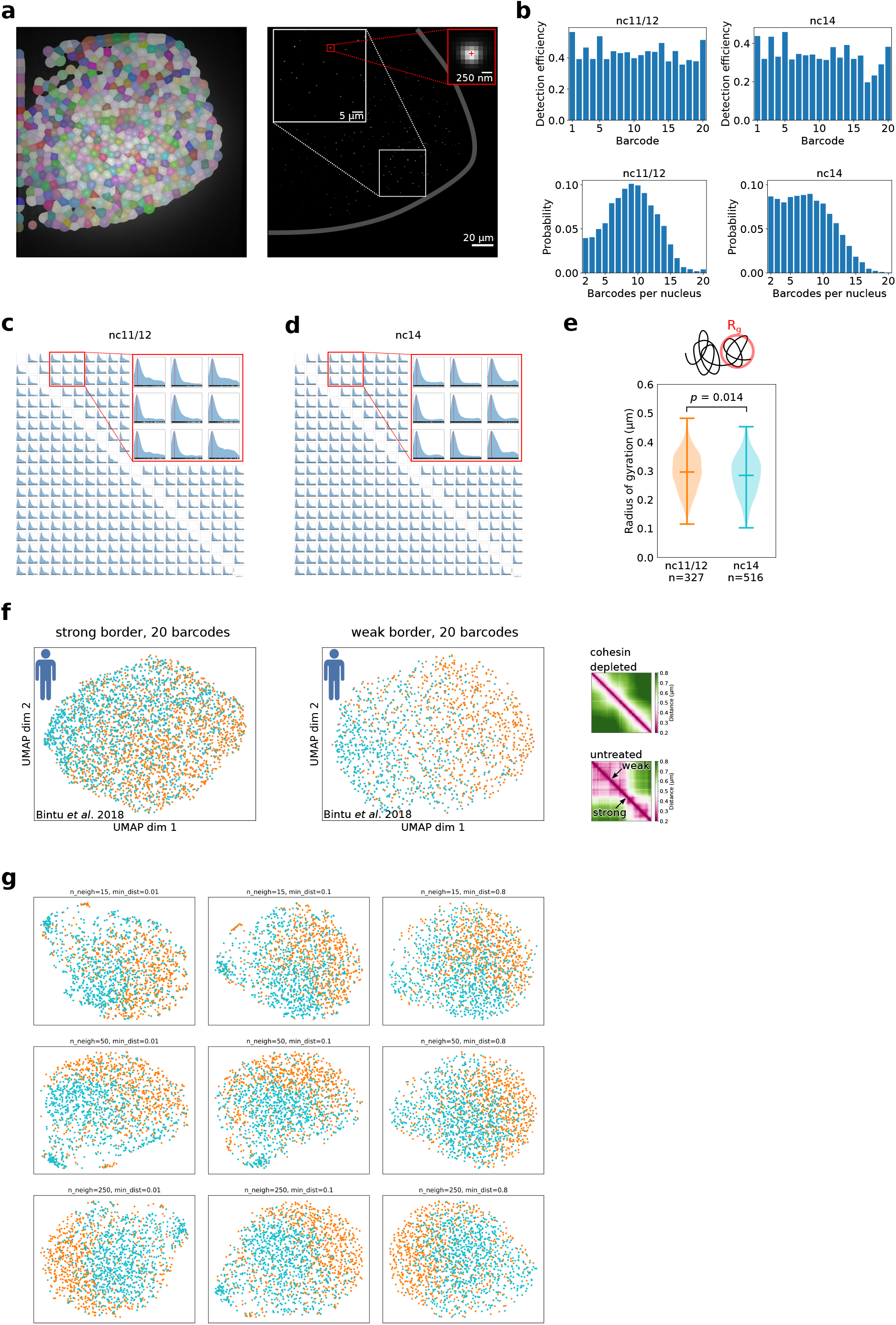
HiM statistics and UMAP validation. **a**. Left: Greyscale image of DAPI-stained nuclei from a *Drosophila* embryo overlaid with the extracted masks after nuclei segmentation. Right: Typical maximum intensity projection of the fluorescence signal from a single barcode in the same field of view as to the left. The border of the embryo is shown by a thick gray line. The red-framed inset shows the point spread function of one barcode and indicates the localization of the center with a red cross. **b**. Top: Detection efficiency for all barcodes, according to nuclear cycle. Bottom: Distribution of the number of detected barcodes per nucleus. **c**. Map of pairwise distance distributions for all barcode combinations in nc11/12 nuclei. The order of the distributions follows that in the ensemble pairwise distance map (Fig. 1b). The blue shade represents a kernel density estimation with a bandwidth of 0.2 μm, the vertical red line represents the maximum of the distribution, and black vertical bars on the x-axis represent individual data points. **d**. Similar to c, but for nc14. **e**. Distribution of the doc-TAD volumes, as measured by its radius of gyration, shown as violin plots. Distribution for nc11/12 in orange (mean Rg=0.30 µm), nc14 in cyan (mean Rg=0.28 µm). Markers indicate the mean and extreme values of the distribution. The *p*-value was calculated by a two-sided Welch’s *t*-test. **f**. Effect of reducing the number of barcodes for the UMAP embedding. Data taken from ^38^. The UMAP on the left was obtained from 20 barcodes centered around the strong TAD border (see black arrows in the ensemble PWD map on the right). The UMAP on the right was obtained from 20 barcodes centered around a weak TAD border. **g**. Segregation of nc11/12 and nc14 nuclei is stable for different UMAP hyperparameters. Plots show UMAPs for different numbers of neighboring sample points (“n_neigh”) and different values for the minimum distance between embedded points (“min_dist”).

**Supplementary Figure 2.**
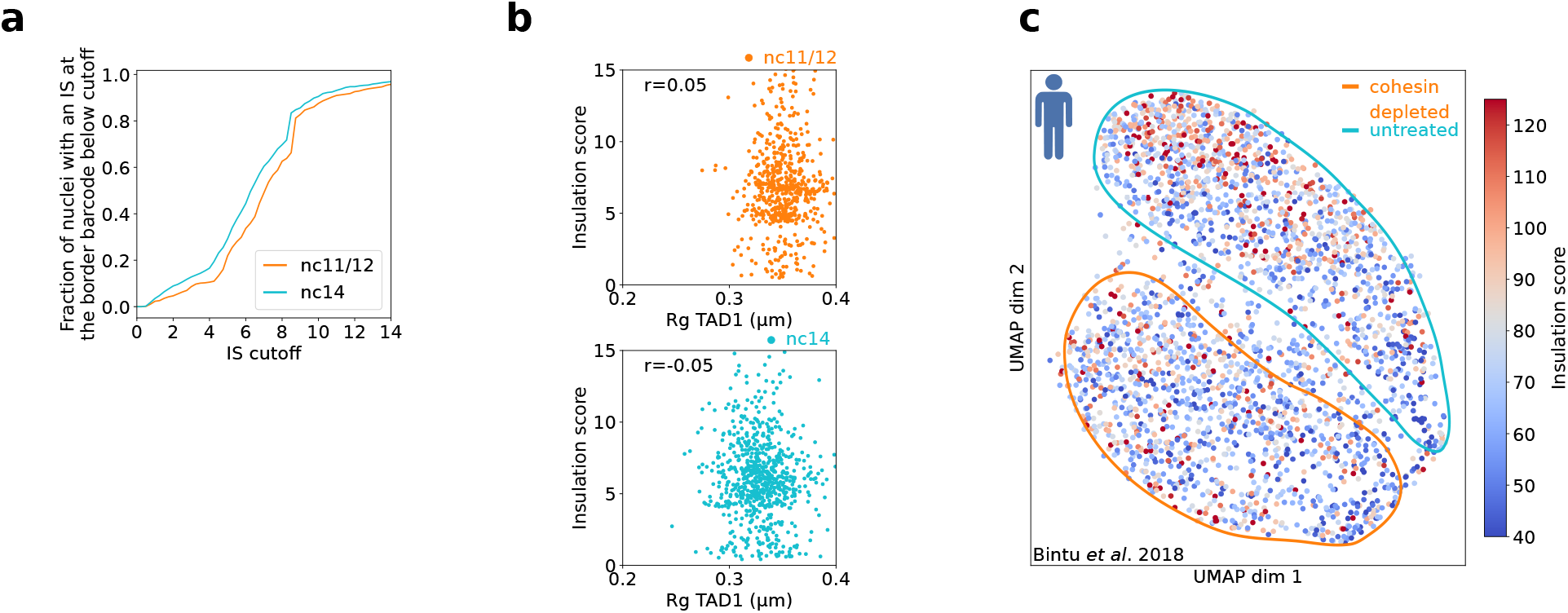
TAD insulation is highly heterogeneous at the single nucleus level. **a**. Cumulative distribution of the single nucleus IS for nc11/12 (orange) and nc14 (cyan) embryos. **b**. Top: scatter plot of insulation score at the border barcode versus radius of gyration of TAD1 in nc11/12. Each point corresponds to a single nucleus. *r* is the Pearson correlation coefficient. Bottom: similar to the top, but for nc14. **c**. Same UMAP as in Fig. 1g (data taken from ^38^), color-coded by the insulation score at the border barcode (indicated as “strong” border in Fig. S1f).

**Supplementary Figure 3.**
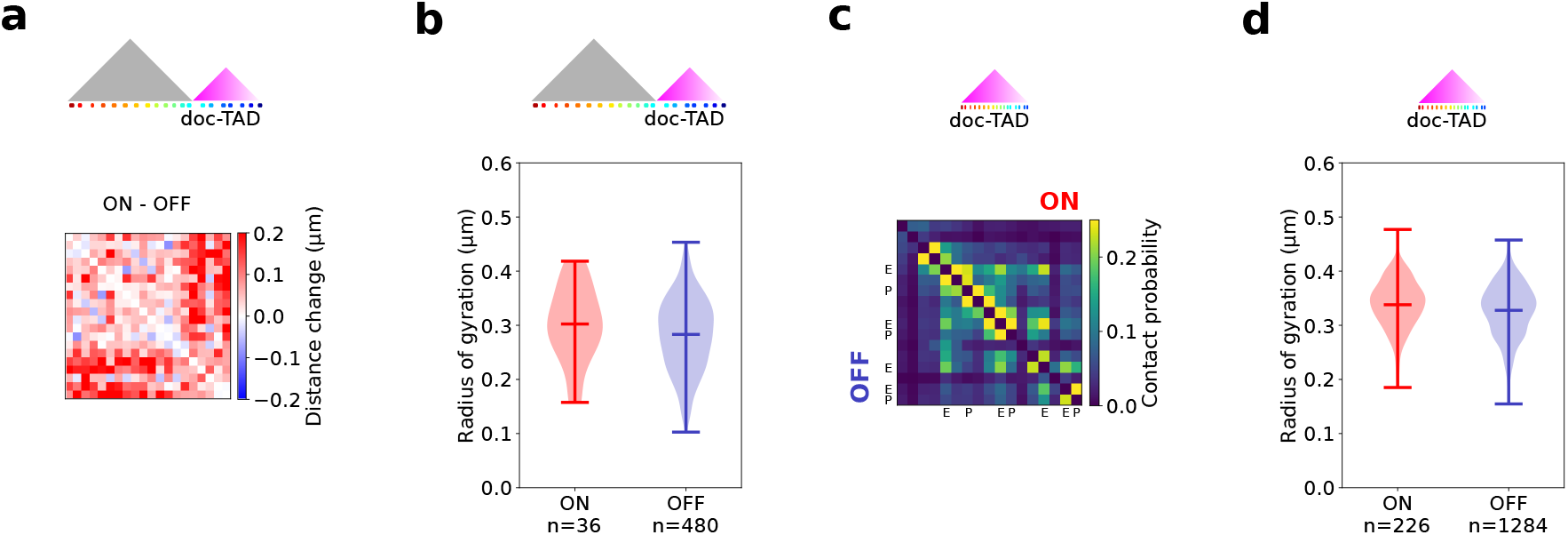
Transcriptional activation has a minor impact on TAD volumes but not proximity frequencies. **a**. Change in the ePWD map between transcriptionally active (“ON”) and inactive (“OFF”) nuclei for the Hi-M library that covers TAD1 and the doc-TAD. Distances that are larger in ON nuclei are in red. **b**. Violin port of the TAD volume (as measured by the radius of gyration) for the doc-TAD in transcriptionally active (“ON”) and inactive (“OFF”) nuclei. Markers indicate the mean and extreme values of the distribution. The radius of gyration is 0.30±0.07µm for active and 0.28±0.07µm for inactive nuclei (mean±standard deviation of the distribution). **c**. Contact probability map for the high-resolution Hi-M library covering the doc-TAD. Upper right half of the matrix displays the map for active nuclei, and the lower left half for inactive nuclei. Barcodes with cis-regulatory elements (enhancer E, promoter P) are indicated. **d**. Similar to panel b, but for the high-resolution Hi-M library. The radius of gyration is 0.34±0.05µm for active and 0.33±0.04µm for inactive nuclei (mean±standard deviation of the distribution).

## References

1. Szabo, Q., Bantignies, F. & Cavalli, G. Principles of genome folding into topologically associating domains. Sci Adv 5, eaaw1668 (2019).

2. Nora, E. P., Lajoie, B. R., Schulz, E. G., Giorgetti, L., Okamoto, I., Servant, N., Piolot, T., van Berkum, N. L., Meisig, J., Sedat, J., Gribnau, J., Barillot, E., Blüthgen, N., Dekker, J. & Heard, E. Spatial partitioning of the regulatory landscape of the X-inactivation centre. Nature 485, 381–385 (2012).

3. Dixon, J. R., Selvaraj, S., Yue, F., Kim, A., Li, Y., Shen, Y., Hu, M., Liu, J. S. & Ren, B. Topological domains in mammalian genomes identified by analysis of chromatin interactions. Nature 485, 376–380 (2012).

4. Sexton, T., Yaffe, E., Kenigsberg, E., Bantignies, F., Leblanc, B., Hoichman, M., Parrinello, H., Tanay, A. & Cavalli, G. Three-dimensional folding and functional organization principles of the Drosophila genome. Cell 148, 458–472 (2012).

5. Banigan, E. J. & Mirny, L. A. Loop extrusion: theory meets single-molecule experiments. Curr. Opin. Cell Biol. 64, 124–138 (2020).

6. Shen, Y., Yue, F., McCleary, D. F., Ye, Z., Edsall, L., Kuan, S., Wagner, U., Dixon, J., Lee, L., Lobanenkov, V. V. & Ren, B. A map of the cis-regulatory sequences in the mouse genome. Nature 488, 116–120 (2012).

7. Symmons, O., Uslu, V. V., Tsujimura, T., Ruf, S., Nassari, S., Schwarzer, W., Ettwiller, L. & Spitz, F. Functional and topological characteristics of mammalian regulatory domains. Genome Res. 24, 390–400 (2014).

8. Neems, D. S., Garza-Gongora, A. G., Smith, E. D. & Kosak, S. T. Topologically associated domains enriched for lineage-specific genes reveal expression-dependent nuclear topologies during myogenesis. Proc. Natl. Acad. Sci. U. S. A. 113, E1691–700 (2016).

9. Ji, X., Dadon, D. B., Powell, B. E., Fan, Z. P., Borges-Rivera, D., Shachar, S., Weintraub, A. S., Hnisz, D., Pegoraro, G., Lee, T. I., Misteli, T., Jaenisch, R. & Young, R. A. 3D Chromosome Regulatory Landscape of Human Pluripotent Cells. Cell Stem Cell 18, 262–275 (2016).

10. Dowen, J. M., Fan, Z. P., Hnisz, D., Ren, G., Abraham, B. J., Zhang, L. N., Weintraub, A. S., Schujiers, J., Lee, T. I., Zhao, K. & Young, R. A. Control of cell identity genes occurs in insulated neighborhoods in mammalian chromosomes. Cell 159, 374–387 (2014).

11. Ron, G., Globerson, Y., Moran, D. & Kaplan, T. Promoter-enhancer interactions identified from Hi-C data using probabilistic models and hierarchical topological domains. Nat. Commun. 8, 2237 (2017).

12. Rao, S. S. P., Huntley, M. H., Durand, N. C., Stamenova, E. K., Bochkov, I. D., Robinson, J. T., Sanborn, A. L., Machol, I., Omer, A. D., Lander, E. S. & Aiden, E. L. A 3D Map of the Human Genome at Kilobase Resolution Reveals Principles of Chromatin Looping. Cell 162, 687–688 (2015).

13. Sexton, T., Yaffe, E., Kenigsberg, E., Bantignies, F. d. R., Leblanc, B., Hoichman, M., Parrinello, H., Tanay, A. & Cavalli, G. Three-dimensional folding and functional organization principles of the Drosophila genome. Cell 148, 458–472 (2012).

14. Hou, C., Li, L., Qin, Z. & Corces, V. Gene density, transcription, and insulators contribute to the partition of the Drosophila genome into physical domains. Mol. Cell 48, 471–484 (2012).

15. Furlong, E. E. M. & Levine, M. Developmental enhancers and chromosome topology. Science 361, 1341–1345 (2018).

16. Bonev, B., Mendelson Cohen, N., Szabo, Q., Fritsch, L., Papadopoulos, G. L., Lubling, Y., Xu, X., Lv, X., Hugnot, J.-P., Tanay, A. & Cavalli, G. Multiscale 3D Genome Rewiring during Mouse Neural Development. Cell 171, 557–572.e24 (2017).

17. Miguel-Escalada, I., Bonàs-Guarch, S., Cebola, I., Ponsa-Cobas, J., Mendieta-Esteban, J., Atla, G., Javierre, B. M., Rolando, D. M. Y., Farabella, I., Morgan, C. C., García-Hurtado, J., Beucher, A., Morán, I., Pasquali, L., Ramos-Rodríguez, M., Appel, E. V. R., Linneberg, A., Gjesing, A. P., Witte, D. R., Pedersen, O., Grarup, N., Ravassard, P., Torrents, D., Mercader, J. M., Piemonti, L., Berney, T., de Koning, E. J. P., Kerr-Conte, J., Pattou, F., Fedko, I. O., Groop, L., Prokopenko, I., Hansen, T., Marti-Renom, M. A., Fraser, P. & Ferrer, J. Human pancreatic islet three-dimensional chromatin architecture provides insights into the genetics of type 2 diabetes. Nat. Genet. 51, 1137–1148 (2019).

18. Schoenfelder, S. & Fraser, P. Long-range enhancer-promoter contacts in gene expression control. Nat. Rev. Genet. 20, 437–455 (2019).

19. Chen, H., Levo, M., Barinov, L., Fujioka, M., Jaynes, J. B. & Gregor, T. Dynamic interplay between enhancer-promoter topology and gene activity. Nat. Genet. 50, 1296–1303 (2018).

20. Benabdallah, N. S., Williamson, I., Illingworth, R. S., Kane, L., Boyle, S., Sengupta, D., Grimes, G. R., Therizols, P. & Bickmore, W. A. Decreased Enhancer-Promoter Proximity Accompanying Enhancer Activation. Mol. Cell 76, 473–484.e7 (2019).

21. Alexander, J. M., Guan, J., Li, B., Maliskova, L., Song, M., Shen, Y., Huang, B., Lomvardas, S. & Weiner, O. D. Live-cell imaging reveals enhancer-dependent transcription in the absence of enhancer proximity. Elife 8, (2019).

22. Ghavi-Helm, Y., Jankowski, A., Meiers, S., Viales, R. R., Korbel, J. O. & Furlong, E. E. M. Highly rearranged chromosomes reveal uncoupling between genome topology and gene expression. Nat. Genet. 51, 1272–1282 (2019).

23. Hug, C. B., Grimaldi, A. G., Kruse, K. & Vaquerizas, J. M. Chromatin Architecture Emerges during Zygotic Genome Activation Independent of Transcription. Cell 169, 216–228.e19 (2017).

24. Ogiyama, Y., Schuettengruber, B., Papadopoulos, G. L., Chang, J.-M. & Cavalli, G. Polycomb-Dependent Chromatin Looping Contributes to Gene Silencing during Drosophila Development. Mol. Cell 71, 73–88.e5 (2018).

25. van der Weide, R. H. & de Wit, E. Developing landscapes: genome architecture during early embryogenesis. Curr. Opin. Genet. Dev. 55, 39–45 (2019).

26. Vallot, A. & Tachibana, K. The emergence of genome architecture and zygotic genome activation. Curr. Opin. Cell Biol. 64, 50–57 (2020).

27. Ghosh, R. P. & Meyer, B. J. Spatial Organization of Chromatin: Emergence of Chromatin Structure During Development. Annu. Rev. Cell Dev. Biol. 37, 199–232 (2021).

28. Ing-Simmons, E., Vaid, R., Bing, X. Y., Levine, M., Mannervik, M. & Vaquerizas, J. M. Independence of chromatin conformation and gene regulation during Drosophila dorsoventral patterning. Nat. Genet. 53, 487–499 (2021).

29. Espinola, S. M., Götz, M., Bellec, M., Messina, O., Fiche, J.-B., Houbron, C., Dejean, M., Reim, I., Cardozo Gizzi, A. M., Lagha, M. & Nollmann, M. Cis-regulatory chromatin loops arise before TADs and gene activation, and are independent of cell fate during early Drosophila development. Nat. Genet. 53, 477–486 (2021).

30. Nagano, T., Lubling, Y., Stevens, T. J., Schoenfelder, S., Yaffe, E., Dean, W., Laue, E. D., Tanay, A. & Fraser, P. Single-cell Hi-C reveals cell-to-cell variability in chromosome structure. Nature 502, 59–64 (2013).

31. Ulianov, S. V., Zakharova, V. V., Galitsyna, A. A., Kos, P. I., Polovnikov, K. E., Flyamer, I. M., Mikhaleva, E. A., Khrameeva, E. E., Germini, D., Logacheva, M. D., Gavrilov, A. A., Gorsky, A. S., Nechaev, S. K., Gelfand, M. S., Vassetzky, Y. S., Chertovich, A. V., Shevelyov, Y. Y. & Razin, S. V. Order and stochasticity in the folding of individual Drosophila genomes. Nat. Commun. 12, 41 (2021).

32. Stevens, T. J., Lando, D., Basu, S., Atkinson, L. P., Cao, Y., Lee, S. F., Leeb, M., Wohlfahrt, K. J., Boucher, W., O’Shaughnessy-Kirwan, A., Cramard, J., Faure, A. J., Ralser, M., Blanco, E., Morey, L., Sansó, M., Palayret, M. G. S., Lehner, B., Di Croce, L., Wutz, A., Hendrich, B., Klenerman, D. & Laue, E. D. 3D structures of individual mammalian genomes studied by single-cell Hi-C. Nature 544, 59–64 (2017).

33. Nagano, T., Lubling, Y., Várnai, C., Dudley, C., Leung, W., Baran, Y., Mendelson Cohen, N., Wingett, S., Fraser, P. & Tanay, A. Cell-cycle dynamics of chromosomal organization at single-cell resolution. Nature 547, 61–67 (2017).

34. Flyamer, I. M., Gassler, J., Imakaev, M., Brandão, H. B., Ulianov, S. V., Abdennur, N., Razin, S. V., Mirny, L. A. & Tachibana-Konwalski, K. Single-nucleus Hi-C reveals unique chromatin reorganization at oocyte-to-zygote transition. Nature 544, 110–114 (2017).

35. Cattoni, D. I., Cardozo Gizzi, A. M., Georgieva, M., Di Stefano, M., Valeri, A., Chamousset, D., Houbron, C., Déjardin, S., Fiche, J.-B., González, I., Chang, J.-M., Sexton, T., Marti-Renom, M. A., Bantignies, F., Cavalli, G. & Nollmann, M. Single-cell absolute contact probability detection reveals chromosomes are organized by multiple low-frequency yet specific interactions. Nat. Commun. 8, 1753 (2017).

36. Finn, E. H., Pegoraro, G., Brandão, H. B., Valton, A.-L., Oomen, M. E., Dekker, J., Mirny, L. & Misteli, T. Extensive Heterogeneity and Intrinsic Variation in Spatial Genome Organization. Cell 176, 1502–1515.e10 (2019).

37. Cardozo Gizzi, A. M., Cattoni, D. I., Fiche, J.-B., Espinola, S. M., Gurgo, J., Messina, O., Houbron, C., Ogiyama, Y., Papadopoulos, G. L., Cavalli, G., Lagha, M. & Nollmann, M. Microscopy-Based Chromosome Conformation Capture Enables Simultaneous Visualization of Genome Organization and Transcription in Intact Organisms. Mol. Cell 74, 212–222.e5 (2019).

38. Bintu, B., Mateo, L. J., Su, J.-H., Sinnott-Armstrong, N. A., Parker, M., Kinrot, S., Yamaya, K., Boettiger, A. N. & Zhuang, X. Super-resolution chromatin tracing reveals domains and cooperative interactions in single cells. Science 362, (2018).

39. Giorgetti, L., Galupa, R., Nora, E. P., Piolot, T., Lam, F., Dekker, J., Tiana, G. & Heard, E. Predictive polymer modeling reveals coupled fluctuations in chromosome conformation and transcription. Cell 157, 950–963 (2014).

40. Jost, D., Carrivain, P., Cavalli, G. & Vaillant, C. Modeling epigenome folding: formation and dynamics of topologically associated chromatin domains. Nucleic Acids Res. 42, 9553–9561 (2014).

41. Conte, M., Fiorillo, L., Bianco, S., Chiariello, A. M., Esposito, A. & Nicodemi, M. Polymer physics indicates chromatin folding variability across single-cells results from state degeneracy in phase separation. Nat. Commun. 11, 3289 (2020).

42. Boettiger, A. N., Bintu, B., Moffitt, J. R., Wang, S., Beliveau, B. J., Fudenberg, G., Imakaev, M., Mirny, L. A., Wu, C.-T. & Zhuang, X. Super-resolution imaging reveals distinct chromatin folding for different epigenetic states. Nature 529, 418–422 (2016).

43. Szabo, Q., Jost, D., Chang, J.-M., Cattoni, D. I., Papadopoulos, G. L., Bonev, B., Sexton, T., Gurgo, J., Jacquier, C., Nollmann, M., Bantignies, F. & Cavalli, G. TADs are 3D structural units of higher-order chromosome organization in Drosophila. Sci Adv 4, eaar8082 (2018).

44. Luppino, J. M., Park, D. S., Nguyen, S. C., Lan, Y., Xu, Z., Yunker, R. & Joyce, E. F. Cohesin promotes stochastic domain intermingling to ensure proper regulation of boundary-proximal genes. Nat. Genet. 52, 840–848 (2020).

45. Szabo, Q., Donjon, A., Jerković, I., Papadopoulos, G. L., Cheutin, T., Bonev, B., Nora, E. P., Bruneau, B. G., Bantignies, F. & Cavalli, G. Regulation of single-cell genome organization into TADs and chromatin nanodomains. Nat. Genet. 52, 1151–1157 (2020).

46. Foe, V. E. & Alberts, B. M. Reversible chromosome condensation induced in Drosophila embryos by anoxia: visualization of interphase nuclear organization. J. Cell Biol. 100, 1623–1636 (1985).

47. Lott, S. E., Villalta, J. E., Schroth, G. P., Luo, S., Tonkin, L. A. & Eisen, M. B. Noncanonical compensation of zygotic X transcription in early Drosophila melanogaster development revealed through single-embryo RNA-seq. PLoS Biol. 9, e1000590 (2011).

48. Farrell, J. A. & O’Farrell, P. H. From egg to gastrula: how the cell cycle is remodeled during the Drosophila mid-blastula transition. Annu. Rev. Genet. 48, 269–294 (2014).

49. Hamm, D. C. & Harrison, M. M. Regulatory principles governing the maternal-to-zygotic transition: insights from Drosophila melanogaster. Open Biol. 8, 180183 (2018).

50. Cardozo Gizzi, A. M., Espinola, S. M., Gurgo, J., Houbron, C., Fiche, J.-B., Cattoni, D. I. & Nollmann, M. Direct and simultaneous observation of transcription and chromosome architecture in single cells with Hi-M. Nat. Protoc. 15, 840–876 (2020).

51. Reim, I., Lee, H.-H. & Frasch, M. The T-box-encoding Dorsocross genes function in amnioserosa development and the patterning of the dorsolateral germ band downstream of Dpp. Development 130, 3187–3204 (2003).

52. Rose, T. The End of Average: How We Succeed in a World That Values Sameness. (HarperCollins, 2016).

53. Sawh, A. N., Shafer, M. E. R., Su, J.-H., Zhuang, X., Wang, S. & Mango, S. E. Lamina-Dependent Stretching and Unconventional Chromosome Compartments in Early C. elegans Embryos. Mol. Cell 78, 96–111.e6 (2020).

54. Crane, E., Bian, Q., McCord, R. P., Lajoie, B. R., Wheeler, B. S., Ralston, E. J., Uzawa, S., Dekker, J. & Meyer, B. J. Condensin-driven remodelling of X chromosome topology during dosage compensation. Nature 523, 240–244 (2015).

55. Nollmann, M., Bennabi, I., Götz, M. & Gregor, T. The Impact of Space and Time on the Functional Output of the Genome. Cold Spring Harb. Perspect. Biol. (2021). doi:10.1101/cshperspect.a040378

56. Nora, E. P., Goloborodko, A., Valton, A.-L., Gibcus, J. H., Uebersohn, A., Abdennur, N., Dekker, J., Mirny, L. A. & Bruneau, B. G. Targeted Degradation of CTCF Decouples Local Insulation of Chromosome Domains from Genomic Compartmentalization. Cell 169, 930–944.e22 (2017).

57. Rao, S. S. P., Huang, S.-C., Glenn St Hilaire, B., Engreitz, J. M., Perez, E. M., Kieffer-Kwon, K.-R., Sanborn, A. L., Johnstone, S. E., Bascom, G. D., Bochkov, I. D., Huang, X., Shamim, M. S., Shin, J., Turner, D., Ye, Z., Omer, A. D., Robinson, J. T., Schlick, T., Bernstein, B. E., Casellas, R., Lander, E. S. & Aiden, E. L. Cohesin Loss Eliminates All Loop Domains. Cell 171, 305–320.e24 (2017).

58. Schwarzer, W., Abdennur, N., Goloborodko, A., Pekowska, A., Fudenberg, G., Loe-Mie, Y., Fonseca, N. A., Huber, W., Haering, C. H., Mirny, L. & Spitz, F. Two independent modes of chromatin organization revealed by cohesin removal. Nature 551, (2017).

59. Chen, X., Ke, Y., Wu, K., Zhao, H., Sun, Y., Gao, L., Liu, Z., Zhang, J., Tao, W., Hou, Z., Liu, H., Liu, J. & Chen, Z.-J. Key role for CTCF in establishing chromatin structure in human embryos. Nature 576, 306–310 (2019).

60. Kaaij, L. J. T., van der Weide, R. H., Ketting, R. F. & de Wit, E. Systemic Loss and Gain of Chromatin Architecture throughout Zebrafish Development. Cell Rep. 24, (2018).

61. Du, Z., Zheng, H., Huang, B., Ma, R., Wu, J., Zhang, X., He, J., Xiang, Y., Wang, Q., Li, Y., Ma, J., Zhang, X., Zhang, K., Wang, Y., Zhang, M. Q., Gao, J., Dixon, J. R., Wang, X., Zeng, J. & Xie, W. Allelic reprogramming of 3D chromatin architecture during early mammalian development. Nature 547, (2017).

62. Buckle, A., Brackley, C. A., Boyle, S., Marenduzzo, D. & Gilbert, N. Polymer Simulations of Heteromorphic Chromatin Predict the 3D Folding of Complex Genomic Loci. Mol. Cell 72, 786–797.e11 (2018).

63. Banigan, E. J., van den Berg, A. A., Brandão, H. B., Marko, J. F. & Mirny, L. A. Chromosome organization by one-sided and two-sided loop extrusion. Elife 9, (2020).

64. Fudenberg, G., Imakaev, M., Lu, C., Goloborodko, A., Abdennur, N. & Mirny, L. A. Formation of Chromosomal Domains by Loop Extrusion. Cell Rep. 15, 2038–2049 (2016).

65. Hansen, A. S., Pustova, I., Cattoglio, C., Tjian, R. & Darzacq, X. CTCF and cohesin regulate chromatin loop stability with distinct dynamics. Elife 6, (2017).

66. Mir, M., Stadler, M. R., Ortiz, S. A., Hannon, C. E., Harrison, M. M., Darzacq, X. & Eisen, M. B. Dynamic multifactor hubs interact transiently with sites of active transcription in embryos. Elife 7, (2018).

67. Dufourt, J., Trullo, A., Hunter, J., Fernandez, C., Lazaro, J., Dejean, M., Morales, L., Nait-Amer, S., Schulz, K. N., Harrison, M. M., Favard, C., Radulescu, O. & Lagha, M. Temporal control of gene expression by the pioneer factor Zelda through transient interactions in hubs. Nat. Commun. 9, 5194 (2018).

68. Coleman, R. A., Liu, Z., Darzacq, X., Tjian, R., Singer, R. H. & Lionnet, T. Imaging Transcription: Past, Present, and Future. Cold Spring Harb. Symp. Quant. Biol. 80, 1–8 (2015).

69. Rajpurkar, A. R., Mateo, L. J., Murphy, S. E. & Boettiger, A. N. Deep learning connects DNA traces to transcription to reveal predictive features beyond enhancer-promoter contact. Nat. Commun. 12, 3423 (2021).

70. Symmons, O. & Raj, A. What’s Luck Got to Do with It: Single Cells, Multiple Fates, and Biological Nondeterminism. Mol. Cell 62, 788–802 (2016).

71. Foe, V. E. & Alberts, B. M. Studies of nuclear and cytoplasmic behaviour during the five mitotic cycles that precede gastrulation in Drosophila embryogenesis. J. Cell Sci. 61, 31–70 (1983).

72. Shaban, H. A., Barth, R., Recoules, L. & Bystricky, K. Hi-D: nanoscale mapping of nuclear dynamics in single living cells. Genome Biol. 21, 1–21 (2020).

73. Dekker, J. & Mirny, L. The 3D Genome as Moderator of Chromosomal Communication. Cell 164, 1110–1121 (2016).

74. Ghosh, S. K. & Jost, D. How epigenome drives chromatin folding and dynamics, insights from efficient coarse-grained models of chromosomes. PLoS Comput. Biol. 14, e1006159 (2018).

75. Popp, A. P., Hettich, J. & Gebhardt, J. C. M. Altering transcription factor binding reveals comprehensive transcriptional kinetics of a basic gene. Nucleic Acids Res. 49, 6249–6266 (2021).

76. Xiao, J. Y., Hafner, A. & Boettiger, A. N. How subtle changes in 3D structure can create large changes in transcription. Elife 10, (2021).

77. Zuin, J., Roth, G., Zhan, Y., Cramard, J., Redolfi, J., Piskadlo, E., Mach, P., Kryzhanovska, M., Tihanyi, G., Kohler, H., Meister, P., Smallwood, S. & Giorgetti, L. Nonlinear control of transcription through enhancer-promoter interactions. (2021). doi:10.1101/2021.04.22.440891

78. ENCODE Project Consortium. An integrated encyclopedia of DNA elements in the human genome. Nature 489, 57–74 (2012).

79. Osterwalder, M., Barozzi, I., Tissières, V., Fukuda-Yuzawa, Y., Mannion, B. J., Afzal, S. Y., Lee, E. A., Zhu, Y., Plajzer-Frick, I., Pickle, C. S., Kato, M., Garvin, T. H., Pham, Q. T., Harrington, A. N., Akiyama, J. A., Afzal, V., Lopez-Rios, J., Dickel, D. E., Visel, A. & Pennacchio, L. A. Enhancer redundancy provides phenotypic robustness in mammalian development. Nature 554, 239–243 (2018).

80. Weigert, M., Schmidt, U., Haase, R., Sugawara, K. & Myers, G. Star-convex polyhedra for 3D object detection and segmentation in microscopy. in 2020 IEEE Winter Conference on Applications of Computer Vision (WACV) (IEEE, 2020). doi:10.1109/wacv45572.2020.9093435

81. Safieddine, A., Coleno, E., Salloum, S., Imbert, A., Traboulsi, A.-M., Kwon, O. S., Lionneton, F., Georget, V., Robert, M.-C., Gostan, T., Lecellier, C.-H., Chouaib, R., Pichon, X., Le Hir, H., Zibara, K., Mueller, F., Walter, T., Peter, M. & Bertrand, E. A choreography of centrosomal mRNAs reveals a conserved localization mechanism involving active polysome transport. Nat. Commun. 12, 1352 (2021).

82. McInnes, L., Healy, J. & Melville, J. UMAP: Uniform Manifold Approximation and Projection for Dimension Reduction. http://arXiv.org (2018). At <https://arxiv.org/abs/1802.03426>

